# A comprehensive experimental comparison between federated and centralized learning

**DOI:** 10.1101/2023.07.26.550615

**Authors:** Swier Garst, Julian Dekker, Marcel Reinders

**Affiliations:** Delft Bioinformatics Lab, Delft University of Technology, van Mourik Broekmanweg 6, Delft, 2628 XE, Zuid-Holland, the Netherlands

**Keywords:** federated learning, distributed machine learning, privacy preserving machine learning

## Abstract

**Purpose:** Federated learning is an upcoming machine learning paradigm which allows data from multiple sources to be used for training of classifiers without the data leaving the source it originally resides. This can be highly valuable for use cases such as medical research, where gathering data at a central location can be quite complicated due to privacy and legal concerns of the data. In such cases, federated learning has the potential to vastly speed up the research cycle. Although federated and central learning have been compared from a theoretical perspective, an extensive experimental comparison of performances and learning behavior still lacks.

**Methods:** We have performed a comprehensive experimental comparison between federated and centralized learning. We evaluated various classifiers on various datasets exploring influences of different sample distributions as well as different class distributions across the clients.

**Results:** The results show similar performances under a wide variety of settings between the federated and central learning strategies. Federated learning is able to deal with various imbalances in the data distributions. It is sensitive to batch effects between different datasets when they coincide with location, similar as with central learning, but this setting might go unobserved more easily.

**Conclusion:** Federated learning seems robust to various challenges such as skewed data distributions, high data dimensionality, multiclass problems and complex models. Taken together, the insights from our comparison gives much promise for applying federated learning as an alternative to sharing data.

## 1 Introduction

Nowadays lots of data is available for the use of machine learning applications. However, in some use cases data does not naturally reside at a single location, and centralizing data might be difficult due to regulation or hardware constraints. One example is medical data ([29], [30]), which is collected at different hospitals or medical institutions, but cannot leave these institutions due to privacy concerns. In order to make use of this type of data, the concept of federated learning ([2]) was introduced. Instead of gathering the data in a centralized location before training a single model, in a federated learning environment, the model gets send to wherever the data is available: the so-called clients. At these clients, the models undergo some form of training, after which the updated model parameters are sent back to a central point, referred to as the server. The server then aggregates all the local updates in order to create a new global model, which it then sends back to all of the clients, and the cycle repeats. With this setup, only model updates are being communicated, and since the original data never has to leave its origins, the entire process becomes less privacy-sensitive.

The concept of federated learning is analogous to other fields trying to learn from distributed data, particularly distributed optimization [13]. Distributed optimization operates similarly to federated learning. The major difference is that federated learning generally uses a star network, with a centralized server connected to each client, whereas such a central point is usually not present in distributed learning, in which clients are only connected to (some) other clients. For a more in-depth analysis of distributed learning we refer the interested reader to [12].

A seminal algorithm for federated learning was Federated Averaging (fedAVG, [2]), where a weighted average of the model parameters is taken at the central server each communication round. Since the introduction of federated learning in 2017, lots of research has been done on its efficiency, performance and privacy-preserving properties [14], [15]. Communication ends up being a bottleneck for efficiency in many federated systems ([31]). As a result, techniques have been developed in order to reduce communication rounds [19], [20]. With regards to performance, many analyses have been made on the performance of federated averaging (fedAVG)[2]. These studies focus usually on performance under poorer data distributions [16], [17], [1], i.e. where data is not independent and identically distributed (IID) at the different clients, as this is the area where fedAVG performance seems to deteriorate [40]. As a result, extensions of federated averaging trying to accommodate for different (non-IID) data distributions have been developed, e.g. [1], [18]. Privacy preservation has been explored by means of constructing specific attacks on federated systems [21], [23]. As a response, extensions on the original federated algorithms that include some form of increased privacy preservation is becoming a vast area of research [22].

Although all of the aforementioned studies have developed the federated learning concept into a vast research area, many analyses remain mostly theoretical. Besides, the federated approach is usually seen as a given, while there are certain use cases where the question whether or not to use federated learning is an important design decision. For example in the medical research field where it is possible, albeit tedious, to gather enough data into a central database. In these cases, one might want to know whether a federated setup will give comparable performance to a central model. Recently, papers comparing a federated setting with centralized baselines have been emerging, e.g. [35], [36]. However, most comparisons only include one or few classifiers, and/or distributions of the data. Therefore, we set out to design a comprehensive set of experiments in which centralized and federated models are compared to one another. We explore the use of different classifiers on multiple datasets, distributed in several ways. In doing so, we shed some light on different scenarios in which federated learning might perform similarly to a centralized model, and when to be careful in assuming such a similar performance.

## 2 Results

### Linear models on a binary classification problem show a relation between the learning rate and amount of clients used

The MNIST dataset (images of handwritten digits) was chosen as afirst dataset, as it is known to be a relatively easy problem for modern day classifiers [5] (methods). In order to simplify, MNIST was converted into a binary classification problem (MNIST2) by selecting only two of the classes. This dataset was distributed evenly (IID) among ten clients, meaning that each client had a similar amount of samples, with an even class distribution (see figure 1(d), methods). A logistic regression (LR) and Support Vector Machine (SVM) (methods) were trained, with varying learning rates for both the central and federated model. Results for the LR model can be found in figure 1(a). These results show that increasing the learning rate leads to faster convergence times, both in the federated and centralized case. There is a consistent factor of ten between the federated and centralized models, i.e. a federated classifier with a learning rate of 0.5 seems to give a comparable result as a central classifier with learning rate 0.05. This was observed for both the LR and SVM models (results on SVM can be found in supplementaryfigure 1). This can be explained by the amount of clients used, which is also ten. A mathematical motivation behind this is given in supplement 1, but the intuition is as follows: a single SGD step in a central model is based on the sum of the loss function over all *S* available data samples. In a federated setting, these *S* datapoints are distributed over *N* clients, meaning that each client gives an update based on only 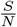 samples (on average). As these updates are only getting averaged, this is analogous to dividing the learning rate by a factor *N*.

**Fig. 1.**
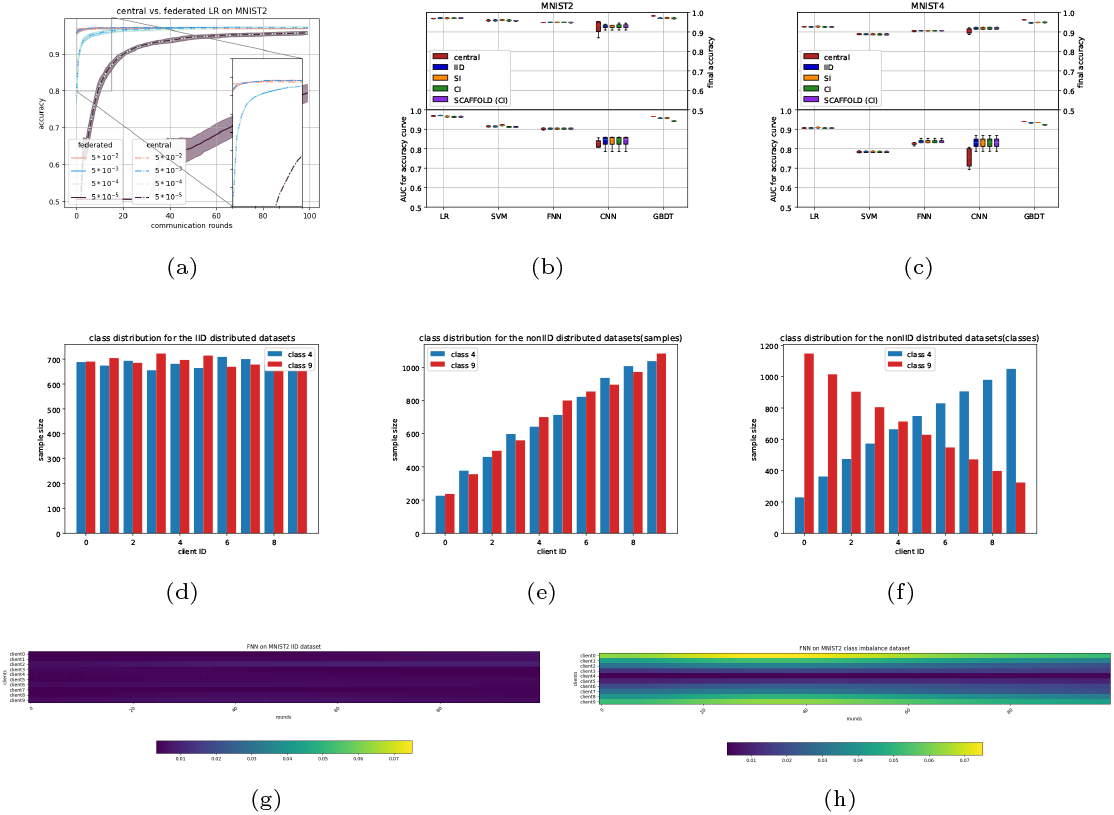
Results on MNIST datasets. (a) shows the relation between federated and centralized learning rate for the Logistical regressor for the MNIST2 dataset. Numbers in the legend correspond to the learning rates. (b)-(c) show a comparison between federated and centralized classifiers for MNIST2 and MNIST4, respectively. (d) - (f) show the various distributions for the MNIST2 dataset (IID, SI, CI, respectively). (g) and (h) show the heatmaps of the euclidean distance between the local and global model parameters after each communication round for the training on the IID and CI distributions, respectively.

### Extension towards more complex models shows different relationships between federated and centralized models

We were interested whether our findings would be different when more complex classifiers were adopted. Hereto, we examined a Fully connected Neural Net (FNN), a Convolutional Neural Net (CNN), and a decision tree (GBDT) classifier (details in methods). Figure 1(b) (supplementary table 4) shows the Area Under Curve (AUC) values for the accuracy curves (i.e. accuracy versus epochs/communication rounds). This metric is indicative of how the convergence of two classifiers compare, especially if the convergence accuracy is equal, which is shown in the top part of the plot. We ran each classifier four times, resulting in the boxpots infigure 1(b). This figure shows that also for the neural network based models (FNN, CNN), similar performance is being reached when comparing the federated (IID) and central models of the same classifier.The GBDT shows a slight difference between the federated and central versions. This could be due to the GBDT using a different federated update scheme as all other classifiers (methods). The CNN shows some difference in AUC of the accuracy curve between the central and federated setting. This highlights a limitation of this metric, as it is caused by a slightly faster start of convergence in the federated case, as can be seen in supplementary figure 2. Furthermore, the convergence accuracy is similar on average, though a higher variance is observed in the central case. Curiously, the relation between learning rate and amount of clients that was observed in the first set of experiments does not seem to be present for the neural network-based classifiers, i.e. these had the same learning rate between federated and central experiments. We speculate that this difference is due to the non-convexity of the neural networks.

### Binary classification experiments show robustness to sample and class distributions

In the next experiment, the robustness to a change in distribution between clients was tested. Hereto, the MNIST2 dataset was split with a sample imbalance across the clients (sample imbalance, SI), i.e. the first client only holds a small fraction of the data, the second client a slightly larger fraction etc. (figure 1(e)). We also considered another imbalance, in which the distribution of classes across clients was skewed (class imbalance, CI) (figure 1(f)), i.e. some clients have more samples from one class and others have more samples from the other class, and in between. Figure 1(b) and supplementary table 4 shows that, with the exception for the GBDT model, similar performances are being attained for the SI and CI settings compared to the IID setting. The SVM does seem to have a slightly higher AUC for the SI distribution, though this is not visible in final accuracy. After observing the accuracy curves (supplementary figure 3), this difference seems to come from a slightly faster convergence rate for the SI distribution (about 1 epoch difference).

Although no performance differences are observed in the imbalance settings, we were interested on how both imbalance settings influenced the learning process. Hereto we visualized the model parameters at the client and at the server at each communication round. In the IID setting (figure 1(g)) one can observe that the updated model parameters at the client are more or less similarly changing across the epochs. Interestingly, for the class imbalance (CI) setting this is not the case, see figure 1(h). Here, we see that the clients with the larger class imbalance have model updates that are consistently further from the global model.

### Extension to multiclass problem results in similar performance as compared to binary classification

To explore the effect of having a multiclass problem, we created a four class dataset out of the original MNIST dataset; MNIST4 (methods). When splitting the samples evenly (IID) across the clients (in number of samples as well as class distributions), the linear models achieve similarly shaped accuracy curves for the federated and central versions, as indicated by the AUC’s in figure 1(c) (supplementary table 4). The neural network based models show differences in these AUCS. In the case of the CNN, this difference is even visible in the final accuracy. Then we distributed MNIST4 with either sample imbalance (SI) or class imbalance (CI), similar to the two class experiments (see supplementary figure 4). The same robustness to the difference in sample and class distributions are observed as in the two class experiments.

### SCAFFOLD model updating performs similar to fedAVG model updating

To be more robust to class imbalances across clients an alternative model updating scheme with respect to fedAVG was introduced: SCAFFOLD ([1], supplementary materials 2). We find that the performance of SCAFFOLD (1(b) and 1(c)) is comparable to fedAVG for the generated distributions of the MNIST2 and MNIST4 datasets. This might have been expected, as fedAVG shows similar performance in the CI case compared to the IID case (so no deteriotation in performance that could be repaired by SCAFFOLD).

### Increasing class distribution aggressiveness gives varying results

After our first experiments on the MNIST datasets, we continued with the fasion MNIST dataset (images of 10 classes of different clothings) as this has been designed to be a harder classification task [5]. When distributing evenly across 01 clients (IID), all models behave similarly as for the more simple MNIST classification tasks (figure 2); there is no visible difference between the central and federated versions of the linear models, the FNN shows a slight difference in AUC but not in final accuracy, and the CNN shows a larger difference in AUC, but not in final accuracy. For the GBDT, some difference can be seen in both AUC and final accuracy.

**Fig. 2.**
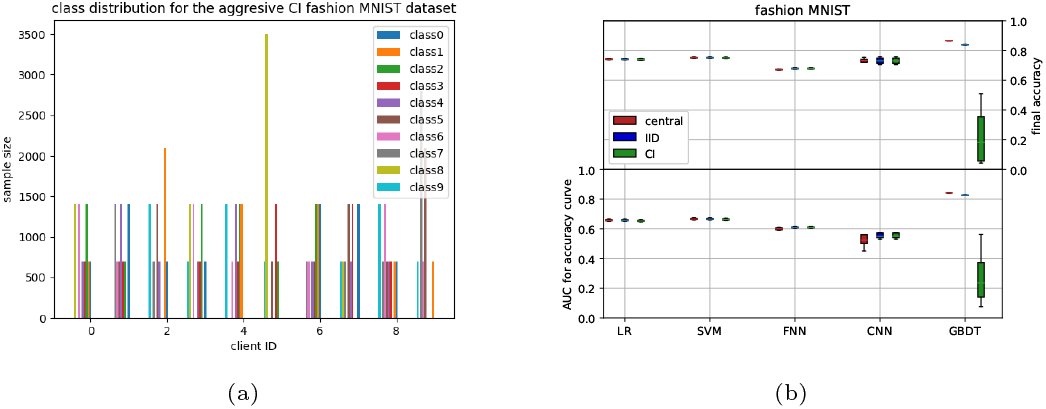
Results on fashion MNIST. (a) shows the data distribution for the aggresive CI distribution for the fashion MNIST dataset. (b) shows results on both IID and CI distributions.

We then tested a more aggressive class imbalance by distributing the fashion MNIST across clients in such a way that the available classes for each client are not necessarily overlapping, i.e. some clients have training examples of a particular class whereas other clients do not (see figure 2(a)). Somewhat surprisingly, differences between the IID setting and the more aggressive CI setting were negligible for most classifiers, except the GBDT, which breaks down for this aggressive CI setting. This might be related to the updating scheme in which the decision tree is updated for each client sequentially (Algorithm 2, methods) instead of simultaneously for all other classifiers (Algorithm 1, methods).

### Experiments on high dimensional data show potential for federated data processing techniques

Until now, all experiments have utilized a single dataset which has then been split into multiple pieces. A more realistic scenario would use multiple datasets, each gathered on a separate client. Hereto, we used datasets that were gathered to predict whether a patient has Acute Myeloid Leukemia (AML) based on their measured transcriptome ([41], methods). It consists of three different datasets (A1-A3), with in total 11.500 patients for which the expression of 12.709 genes are measured. For two of the datasets (A1, A2) the expresssion is measured using microarray technology, and for the third dataset (A3) this was done using current RNA sequencing technology.

In order to determine the feasibility of learning on these datasets, we first analysed A2 only. A2 was distributed in an IID fashion over ten clients. As this data has a high dimensionality (over 12.000 features), a dimensionality reduction method seemed to be essential as all classifiers could not perform above random level in the original feature domain (this was also the case for the central models). Therefore, a Principal Component Analysis (PCA) was performed beforehand, reducing the data to 100 features. For this we implemented the PCA in a federated manner (adapted from [34], see supplementary algorithm 3). With the reduced dataset, all classifiers perform similarly to what we observed in earlier experiments (figure 3(a)). Next, we also tested the SI and CI distributions (before reapplying the federated PCA). Also for this distributed A2 dataset we observe a high robustness to the different data distributions, see figure 3(a) and supplementary table 4.

**Fig. 3.**
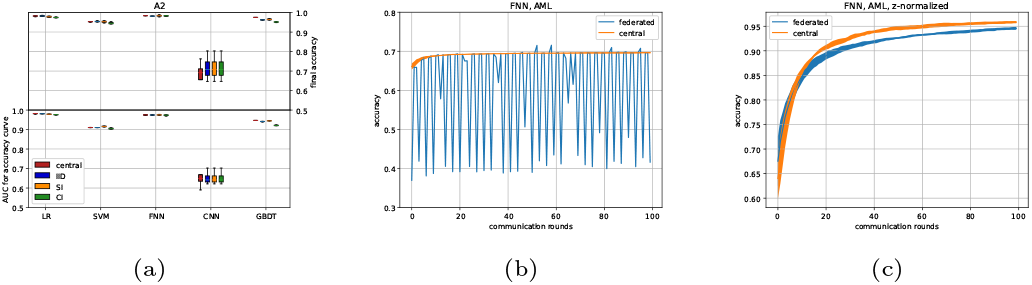
Results on the AML datasets. (a) AUC performance of the four classifers across different splits of dataset A2 across ten clients, (b) FNN results when all three AML datasets A1-A3 are used with the original data. (c) FNN results on A1-3 using the locally scaled data.

### Usage of heterogeneous datasets highlights the need to ‘think federated’

Then we returned to the combined datasets A1-A3. In this setup, only three clients have been used, each holding one complete dataset. Also in this setting dimensionality reduction using federated PCA was done before training the classifiers. The Logistic regressor seems to perform fine, although the relation between client amount and learning rate seems to have disappeared (supplementary figure 5). However, whereas the centralized neural network manages to improve slightly over time, the federated neural network now breaks down. (see figure 3(b)). We therefore dived deeper into possible causes of this breakdown. This breakdown might have been caused by difference in measuring technology, causing the measured features to be distributed differently between dataset A3 (sequencing technology) and the datasets A1 and A2 (microarray technology). To correct for the different feature distributions between the datasets, we normalised features for all datasets locally, similar to a batch correction scheme, by z-normalizing each gene before applying the federated PCA (supplementary algorithm 2). After this batch correction, performance of the federated and central classifiers are similar again (figure 3(c)). These results underline the importance of treating distributed datasets as separate entities, instead of as subsets of a larger dataset.

### Experiments on ‘real life’ distributed datasets show increased variance for federated models

Next, we explored a dataset consisting of molecules that are described by their molecular fingerprints (deMorgan fingerprints, see methods for details) which can be used to predict the activation or inhibition of certain kinases [38]. The dataset is compiled out of three different studies, which we treated as different clients. Two different kinases, KDR and ABL1, were selected as targets for classification, resulting in two datasets. Note that, due to sparsity in the data, these two datasets are not equal in size (for each client), see figures 4(a) and 4(d) for data distributions. After a brief analysis of the datasets, two interesting aspects came to mind. First, the distributions shown in figure 4 a and d could be seen as a combination between a class -and sample imbalanced dataset, thereby increasing complexity w.r.t. earlier distributions. Second, we looked at a combined tSNE plot (figures 4(b) and 4(e)), which show a large amount of small clusters, most of which are only present at one of the datasets.

**Fig. 4.**
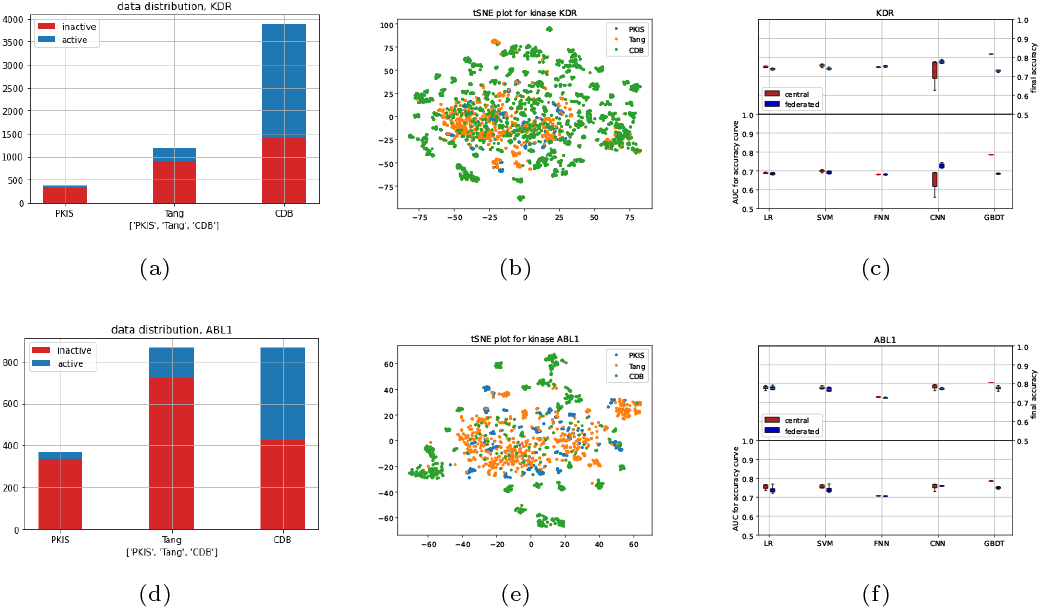
Results on kinase dataset for two kinases, KDR and ABL1. (a) and (d) show data distributions, (b) and (e) tSNE plots (every dot is a molecule, color indicates study), and (c) and (f) show performance results for the different classifiers for the central and federated setting.

Figures 4(c) and 4(f) show the results for kinase KDR and ABL1, respectively, as well as supplementary table 1. For KDR, convergence behavior seems similar, albeit with a higher variance for the linear models, with the exception of CNN and GBDT. However, some difference in final accuracy can be seen for the linear models as well. CNN seems to outperform on KDR with the given metric, which can also somewhat be observed in Supplementary figure 6. For this experiment, minibatch learning was used for the CNN experiments (both federated and centralized), which could explain this behaviour.

For ABL1, there is a larger difference in convergence for the linear models between the federated and centralized settings. However, final accuracy is still similar, as can also be seen in supplementary figure 7. Curiously, both FNN and CNN seem to converge similarly this time, though there is a slightly higher final accuracy for the CNN.

### Complex federated dataset shows capability for larger neural networks

To compare performances on a more complex, distributed dataset, we used the publicly available dataset for the MNM challenge from [37]. 3d MRI scans of the heart are classified as either a healthy patient, or as a patient having one of various cardiovascular diseases. We selected only MRI scans of healthy patients and patients with hypertrophic cardiomyopathy (HCM), which was the largest class of disease, simplifying the task to a binary classification.

After initial testing, it became clear that the structure of the 3d MRI scans was too complex for the simple linear and shallow neural networks to achieve reasonable performances. We therefore ran ResNet-34 with a custom classification head (see methods). For the federated experiment, the dataset was split based on hospital of origin. This resulted in four clients (the fifth hospital did not have any samples of the classes we selected), see figure 5(a). The train/test split chosen by the original authors proved to be exceptionally challenging without any data augmentation strategies, as performed in earlier work on the same dataset [37] [39]. We therefore changed the train/test split as shown in figure 5(b) (note that data stayed at its original ‘location’, only local changes to the train/test splits were made). Figure 5(c) shows the comparison between the federated and centralized experiments. This shows, although the learning behaviour between the two settings is different, both the central and federated approach converge to a similar accuracy, indicating that federated learning also works well on larger (neural) networks, which is in line with earlier research [39].

**Fig. 5.**
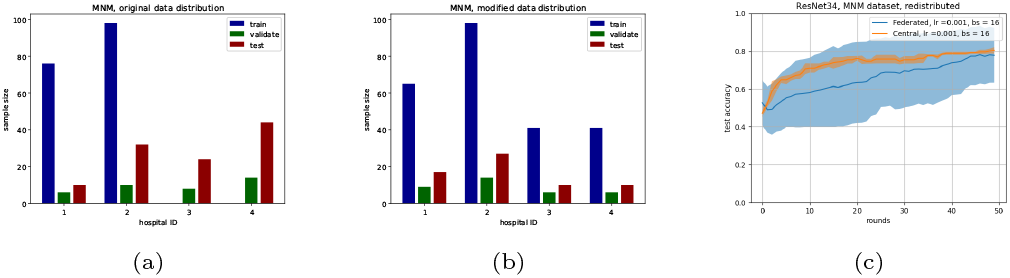
Results on the MNM dataset. (a) and (b) show distributions of the MNM dataset, originally and after redistribution, respectively. (c) shows the acccuracy curve of ResNet 34 on the distribution from (b) (bs = batch size).

## 3 Discussion

This paper describes the results of a set of experiments exploring the differences between centralized and federated machine learning models. Multiple classifiers were used on several datasets, which have been distributed in different ways. Results show that, especially under IID circumstances, a federated classifier could reach similar performances compared to its centralized counterpart. However, a careful choice of parameters such as learning rate and batch size is vital to reach comparable performance. The exact relation between those parameters for reaching equal performance between centralized and federated models remains unclear and could be part of future work, although we have shown that the learning rate can be adapted taking into account the number of clients. Furthermore we have illustrated that when using datasets from multiple sources, one can suffer from batch effects between the datasets. Although this affects both a federated as well as a central model, we advocate to be extra careful on this aspect in the federated case as the data distributions are not trivially compared in this setting.

The federated GBDT-classifier breaks down on the aggresive class-imbalanced distribution of the fashion MNIST dataset, in which each client has approximately half of the total amount of classes. The inPrivate learning algorithm utilizes decision trees created by the previous clients to boost. If these trees were created using a different subset of classes, it can be understood that the resulting gradient does not serve as a helpful tool for the creation of the next decision tree.

For the AML dataset we have shown the necessity of dimension reduction to achieve reasonable performance for the classifiers studied. We should note that in the original work [11], no form of dimension reduction was used. This might have been possible because they used a deep neural network including multiple dropout layers, which are known for being helpful at dealing with highly dimensional data ([33]). In contrast, the neural network used in this paper only consisted of two linear layers, combined with a relu-layer.

One of the strengths of the federated averaging algorithm is that it allows for multiple local epochs, as well as batch learning. This results in multiple learning steps per communication round, potentially reducing the total amount of communication rounds required [2]. In our work, we did not explore the amount of local epochs or batch learning in the federated setting. A further analysis on differences between federated and central models could include experiments exploring the influence of varying these parameters.

One limitation of the results on all MNIST datasets is that it is composed of one original dataset. This means that there are no concept shifts (i.e. batch effects) between the different clients, even in the class -and sample imbalanced cases, as can be observed in the AML dataset. Another limitation is the lack of calibration. Calibration is especially interesting in a federated learning case, as there can be a high variance in class distribution between clients (as has been simulated in this work). How this influences a federated classifier is an interesting direction for future work.

Hospitals could decide to ‘personalize’ their model, i.e. take a model trained using a federated approach (with or without their local dataset), to then retrain that model on their local data only. Although this might lose some generalizability, it could also result in a more accurate model for the institution itself. Exploring such strategies is an interesting venue for future work as well.

Taken together, our work has shown promising results for federated learning as an alternative to centrally organised learning. We strongly advice that, like in central learning with batch effects, it is important to align data distributions across the different clients. Nevertheless, federated learning might open new possibilities to increase data availability for data hungry machine learning methods.

## 4 Methods

### 4.1 Algorithm overview

In total, five different classifiers were implemented in a federated setting: a Logistic Regressor (LR), a Support Vector Machine (SVM), a Fully connected Neural Network (FNN), a Convolutional Neural Network (CNN) and a Gradient-Boosting Decision Tree protocol (GBDT). All classifiers except the GBDT follow the same federated learning scheme, which is given by Algorithm 1. The federated learning algorithm for the GBDT is given by Algorithm 2.

The architectures for the FNN and CNN can be found in supplementary tables 2 and 3 respectively. A performance comparison with a centralized version of the classifiers was the outcome of interest, rather than optimizing the final performance. Therefore the neural net architectures were kept simple, allowing for faster training. The linear models were implemented using sklearn’s SGDClassifier, using default parameters (except for the loss function). Similarly, the GBDT protocol was implemented using sklearn’s GradientBoostingClassifier, using default parameters except for *n estimators*, which was initialized as one, and incremented each iteration to allow for the construction of subsequent trees.

Algorithm 1 works as follows: First, a model is initialized on the server with random values. This model is then sent to all clients. Upon reception, every client performs (in parallel) a local training step to update the coefficients of the local model based on Stochastic Gradient Descent (SGD). All clients then send back their updated model to the central server. At the server, these models are combined to update the global model. Then the next epoch starts, and the combined coefficients from the previous round are now used as the new values to be sent out to all clients. In Algorithm 1, the intermediate models are evaluated on the test set at the server (holding a concatenation of all test sets from all clients). Although this is not realistic, we observed no difference between calculating the performance locally at each client and then averaging the performances, or doing this centrally on the combined test sets. Therefore, for simplicity reasons, we chose to calculate the performances centrally.

#### Algorithm 1 The federated learning algorithm of all classifiers except GBDT

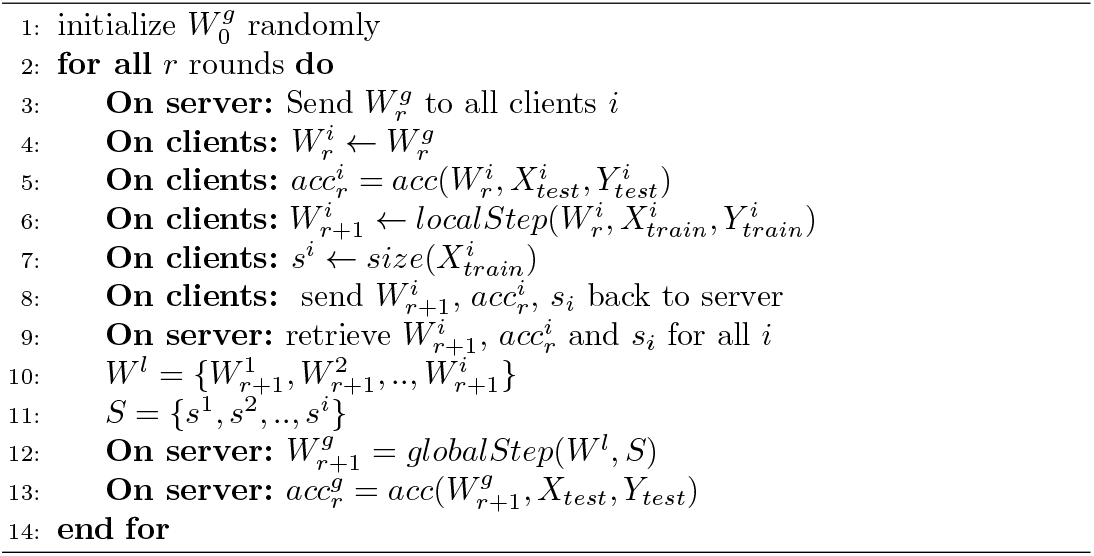

The functions *localStep* and *globalStep* in lines 6 and 12 of algorithm 1 respectively, are either an implementation of the federated averaging Algorithm ([2]), or the SCAFFOLD algorithm ([1]).

The *localStep* for federated averaging is done by means of Stochastic Gradient Descent, i.e.:

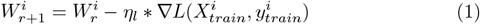

where 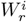 is the model on client *i* in round *r, η*_*l*_ the local learning rate, 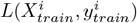 is a loss function which differs between classifiers, and 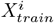 and 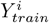 are the training data available at client *i*. Note that the clients also have to split data into training and testing to be able to independently evaluate a learned classifier. The global step of federated averaging consists of taking the weighted average for all model parameters:

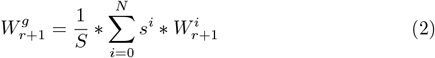

where *s*^*i*^ is the dataset size of the *i*^th^ client, *N* the amount of clients and 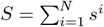.

Besides federated averaging, we have also used the SCAFFOLD aggregation algorithm [1]. For both the local and global step, SCAFFOLD expands on federated averaging by means of a so-called control variate *c*. Intuitively, these control variates are used to compensate for model drift, where a model update does not move into the direction of the global optimum due to local datasets being distributed in a non-IID fashion. For the local step:

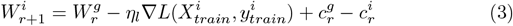

in which 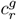 and 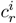 are the global and local control variates in round *r* which are all initialized with zero values. Next (but within the same round), *c*^*i*^ gets updated as:

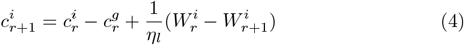

These updated control variates are sent back to the server together with the model parameters and accuracy for that round. Then the global update step of SCAFFOLD first updates the coefficients:

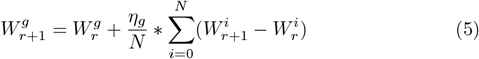

with *η*_*g*_ being the global learning rate. The global control variate *c*^*g*^ :

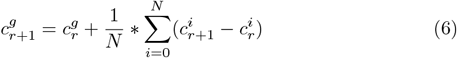

The *localStep* of both fedAVG and SCAFFOLD allows for batch learning, i.e. splitting up the local training data into multiple shards, allowing multiple consecutive gradient descent steps each local epoch. However, our main objective was comparison with the central case and not optimizing performance. Therefore, since batch learning was not required for convergence, it was not utilized. Exception to this is the CNN classifier for the kinase datasets, which needed some batch learning for convergence: For all CNN experiments, the local data was split up into 10 batches each epoch.

#### Algorithm 2 The inPrivate learning algorithm

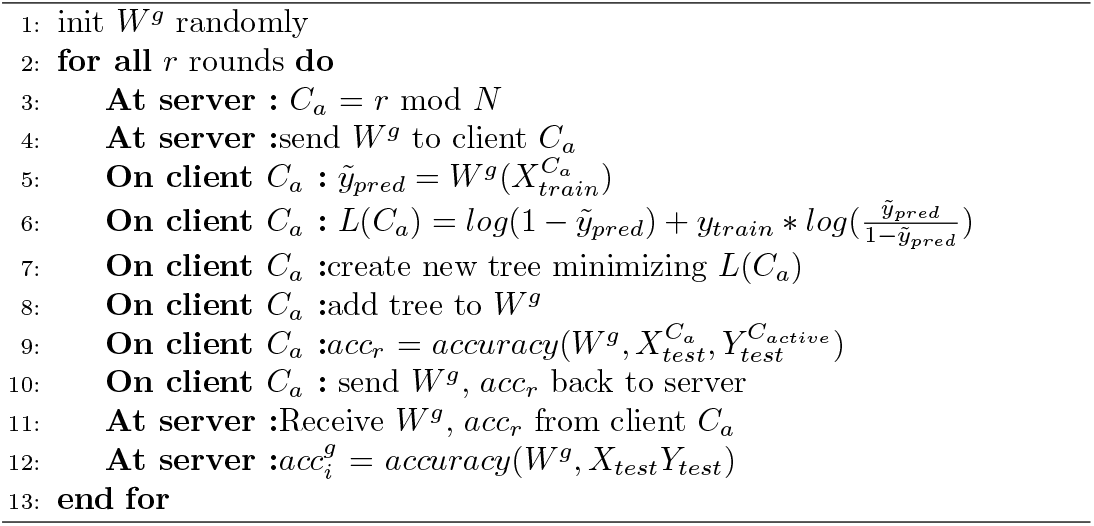

The federated GBDT is implemented differently from the other classifiers, as a Decision Tree does not lend itself very well for parameter averaging. Instead, for GBDT, we followed the “inPrivate Learning” algorithm from ([4]), as shown in Algorithm 2. In this algorithm, only one client is active per round. This client is referred to as *C*_*a*_ in Algorithm 2. This client uses the decision trees of its predecessors to calculate a loss on its own dataset, which it then uses to boost the decision tree it builds.

### 4.2 Datasets

All classifiers were tested on five different datasets: i) MNIST [6], images of handwritten digits, ii) fashion MNIST [5], images of clothes, iii) a dataset used for the prediction of Acute Myeloid Leukemia (AML) based on measured gene expressions [11], iv) a dataset constisting of molecular fingerprints used to predict the activation of certain kinases, and v) a dataset of 3d MRI scans of hearts from either healthy patients or patient with a cardiovascular disease.

#### Two and Four Class MNIST

The first dataset is derived from MNIST [6]. In order to reduce complexity, two derivatives were made, using only a subset of the original classes: a two-class problem (MNIST2) and a four-class problem (MNIST4). The digits were chosen based on which combination is most difficult to separate according to the original work introducing MNIST. For MNIST2, we included digits 4 and 9, whereas for MNIST4, digits 2 and 8 are being added. 80% of the dataset is used for training, with the remaining 20% for testing. Three different splits were made, resulting in 6 federated datasets: an IID distribution, a distribution with imbalances in sample size (SI) and one with imbalances in classes (CI). Figure 1(d) through 1(f) and supplementary figure 4 show these distributions.

#### Fashion MNIST

The fashion MNIST dataset [5] was also split into 80% training and 20% test data. Two different distributions were made: an IID distribution and a class-imbalanced distribution. Note that this imbalanced distribution is different from the MNIST distributions as it concerns ten classes, see figure 2(a).

#### AML dataset (A1-A3)

The third dataset was taken from a study done by Warnat-Herresthal et al. [11]. In their experiments, they used measured transcriptomes of gene expressions from patients to predict the presence of acute myeloid leukaemia (AML). These transcriptomes were taken from human peripheral blood mononuclear cells (PBMC) from three different sources. In the datasets, A1 and A2 are transcriptomes that have been measured using microarrays ([24]), whereas dataset A3 is measured using RNA-sequencing ([25]). All three datasets contain 12709 different gene expression values per sample, with labels for 25 different illnesses. Samples were labelled according to ([11]), i.e. all AML samples were given label (1), and all other labels were joined under one label (0) (in the original work named ‘cases’ and ‘controls’). This approach does result in an inherent class imbalance in the combined dataset, i.e. 7500 samples have label 0 and only 4000 samples have label 1. The datasets were split into 80% training samples and 20% test samples.

#### Kinase dataset

The kinase dataset was taken from [38]. It consists of two deMorgan fingerprints, connectivity and feature based, with as labels the activation values for certain kinases. We chose two of these kinases, being KDR and ABL1. Two of the three datasets were quite sparse, so there is no full overlap between molecules with data on KDR and ABL1, practically creating two datasets. The activation values were binarized between active and non-active, with a cutoff value of 6.3 (e.g. a molecule is seen as activating a kinase if it has a *pIC*_50_ value higher than 6.3), following the original study.

#### MRI dataset

The last dataset consisted of 3d MRI scans of patients’ hearts, taken from [37], where it was generated as part of a challenge on using deep learning models for cardiac MRI scans. Multiple time frames of MRI scans are available for all patients’. Following instructions from [37], only the time points of the end-diastolic and the end-systolic phase were used. As various MRI scanners were used (at different hospitals), not all scans had the same resolution/dimensionality. Scans were brought to equal dimensionality by downsampling to the lowest common dimensions.

We only selected samples from healthy patients and patients with hypertrophic cardiomyopathy (HCM), making it a binary classification problem. The original dataset had a train/validate/test split set up to make the problem extra challenging. Since our main objective is not to solve this issue, but rather compare federated and central classifiers, we opted to redistribute the test/validate/train split to be more similar across all clients, see figure 5(b). Note that this did not mix the samples across clients.

### 4.3 Experimental setup

Our goal is to compare a federated classifier performance to a central model, i.e. when all data is available at the server. To do this in a fair way, the learning rate was kept equal between the federated and central runs. Also, the amount of communication rounds was kept equal to the amount of epochs in the central case. Only one ‘local’ epoch was used during all experiments, meaning all classifiers sent their updated model back to the server after only one full pass of their local data. Four runs with different model initializations were made per experiment (for both the federated and centralized case). All experiments were executed on one laptop, running all clients as well as the server. The federated learning platform vantage6 [9], [10] was used. Vantage6 uses a dockerized solution, which means that every client (and every task that every node runs) runs in its own docker, therefore being unable to alter the solution of the other clients. The experiment on the MRI dataset, however, was not performed with vantage6 due to size constraints of both data and neural network. For this case, we built a simulated federated environment to run on our high performance cluster.

## Data availability and accessability

Code to reproduce the findings of the work can be found here: https://github.com/swiergarst/FLComparison

## Notes

### Competing Interest Statement

The authors have declared no competing interest.

