## Supplementary Material for "A comprehensive experimental comparison between federated and centralized learning"

#### 1 Relation between clients and learning rate (for linear models)

The local model update from federated averaging (as used during this paper) is given by stochastic gradient descent:

$$W_t^i = W_{t-1}^g - \eta \nabla L(W_{t-1}^g(X^i), y^i) \tag{7}$$

These local model updates are then combined into a new global model as follows:

$$W_t^g = \frac{1}{S} \sum_{i=1}^{N} s^i W_t^i \tag{8}$$

Where $N$ is the amount of clients, $s^i$ the dataset size of client $i$, and $S = \sum_{i=1}^{N} s^i$. If we combine equations 7 and 8, we get:

$$W_t^g = \frac{1}{S} \sum_{i=1}^{N} \{s^i [W_{t-1}^g - \eta \nabla L(W_{t-1}^g(X^i), y^i)]\}. \tag{9}$$

By doing some refactoring, equation 9 becomes:

$$W_t^g = \frac{1}{S} \sum_{i=1}^{N} s^i W_{t-1}^g - \frac{1}{S} \sum_{i=0}^{n} s^i \eta \nabla L[W_{t-1}^g(X^i, y^i)] = \frac{W_{t-1}^g}{S} \sum_{i=1}^{n} s^i - \frac{1}{S} \sum_{i=1}^{n} s^i \eta \nabla L[W_{t-1}^g(X^i), \tag{10}$$

Since $S = \sum_{i=1}^{n} s^i$, we get:

$$W_t^g = W_{t-1}^g - \frac{1}{S} \sum_{i=0}^{N} s^i \eta \nabla L[W_{t-1}^g(X^i), y^i]. \tag{11}$$

The loss function can be written as a sum of the loss over all samples in the local dataset, i.e. :

$$L(W_t^g(X^i), y^i) = \sum_{j=0}^{s^i} L(W_t^g(x_j^i), y_j^i), \tag{12}$$

with $x_j^i$ the $j^{th}$ sample in dataset $X^i$, and $y_j^i$ its corresponding label. Combining equation 11 and 12, we get:

$$W_t^g = W_{t-1}^g - \frac{1}{S} \sum_{i=0}^{N} s^i \eta \nabla \sum_{j=0}^{s^i} L(W_t^g(x_j^i), y_j^i) = W_{t-1}^g - \frac{1}{S} \sum_{i=0}^{N} s^i \eta \nabla \sum_{j=0}^{s^i} L_j^i \tag{13}$$

where $L_j^i$ is used as shorthand for $L(W_t^g(x_j^i), y_j^i)$. From here we can work towards showing the relation between the amount of clients and learning rate. There are two separate assumptions that can be made, which lead to similar analyses with the same result. Both analyses will be given. The options are:

1. Assume that each client holds an approximately equal amount of samples, i.e. $s^1 = s^2 = ... = s^i$ for each $i \in N$
2. Assume that each client gives an equivalent model update each round, i.e. $\nabla L_r^1 = \nabla L_r^2 = ... = \nabla L_r^i$ for each $i \in N$

Note that the second assumption is easiest to statisfy if the samples are IID distributed. However, although unlikely, a scenario is possible in which one clients holds only a few samples which result in a large loss each, whereas most clients hold many more samples with only a small loss per sample. Therefore, it is not strictly necessary to assume IID distribution with assumption 2.

#### 1.1 Using assumption 1

With assumption 1, we can rewrite $s^i$ as:

$$s^1 = s^2 = ... = s^i = \frac{S}{N}. \tag{14}$$

Now, using equation 14 with equation 13, we get:

$$W_t^g = W_{t-1}^g - \frac{1}{S}\sum_{i=0}^{N}\frac{S}{N}\eta\nabla\sum_{j=0}^{s^i}L(W_t^g(x_j^i)y_j^i) = W_{t-1}^g - \frac{\eta}{N}\nabla\sum_{i=0}^{N}\sum_{j=0}^{s^i}L(W_t^g(x_j^i), y_j^i). \tag{15}$$

In order to compare with a centralized case, lets assume a setting in which one single client holds all $S$ datapoints. a similar (no batch learning) SGD step of such a client can be described as:

$$W_t^c = W_{t-1} - \eta\nabla L(W_{t-1}(X), y), \tag{16}$$

where $W_t^c$ is the central model at epoch $t$, and

$$X = \sum_{i=1}^{N} X^i, \tag{17}$$

is the concatenation of all local datasets $X^i$. Combining 16 and 17, we get:

$$W_t^c = W_{t-1}^c - \eta\nabla\sum_{i=1}^{N} L(W_{t-1}^c(X^i), y). \tag{18}$$

Now using equation 12, we can get:

$$W_t^c = W_{t-1}^c - \eta \nabla \sum_{i=1}^{N} \sum_{j=1}^{s^i} L(W_{t-1}^c(x_j^i), y). \tag{19}$$

Comparing equations 15 and 19, it becomes clear that they are equivalent, except for the factor $\frac{1}{N}$ in equation 15, which effectively lowers the learning rate of the federated case by a factor $N$ as compared to its central counterpart.

##### 1.2 Using assumption 2

For this analysis, it is convenient to use equation 11 as a starting point instead of equation 13:

$$W_t^g = W_{t-1}^g - \frac{1}{S} \sum_{i=0}^{N} s^i \eta \nabla L^i, \tag{20}$$

where we use $L^i$ as shorthand for $L(W_{t-1}^g(X^i), y^i)$. Since we assume that $\nabla L^1 = \nabla L^2 = \nabla L^i$ for all $i \in N$, we can remove the dependency of i, i.e.

$$W_t^g = W_{t-1}^g - \frac{1}{S} \sum_{i=0}^{N} s^i \eta \nabla L^i = W_{t-1}^g - \frac{1}{S} \sum_{i=0}^{N} s^i \eta \nabla L. \tag{21}$$

Since $S = \sum_{i=0}^{N} s^i$, we now get:

$$W_t^g = W_{t-1}^g - \frac{1}{\sum_{i=0}^{N} s^i} \sum_{i=0}^{N} s^i \eta \nabla L = W_{t-1}^g - \eta \nabla L. \tag{22}$$

Comparing to the central case, we start off with equation 19:

$$W_t^c = W_{t-1}^c - \eta \nabla \sum_{i=1}^{N} \sum_{j=1}^{s^i} L(W_{t-1}^c(x_j^i), y) = W_{t-1}^c - \eta \nabla \sum_{i=1}^{N} L^i, \tag{23}$$

where we once again use $L^i = L(W_{t-1}^g(X^i), y^i)$. Once again, using $L^i = L$, we arrive at:

$$W_t^c = W_{t-1}^c - \eta \nabla \sum_{i=1}^{N} L^i = W_{t-1}^c - \eta \nabla \sum_{i=1}^{N} L = W_{t-1}^c - N \eta \nabla L. \tag{24}$$

#### 2 SCAFFOLD

The SCAFFOLD algorithm was created as a means to increase the performance of the earlier discussed federated averaging, specifically with respect to a non-IID setting. A non-IID setting means that the distribution of classes between different clients is not similar, i.e. some clients only having access to a very low amount (in some cases even zero) of samples of certain classes. The problem that arises in federated averaging within the non-IID setting can be intuitively described as a drift, meaning that the global parameters do not or slowly converge to the optimum. In order to combat this drift, [1] introduce a new algorithm for Stochastic Controlled Averaging, called SCAFFOLD.

The main addition in SCAFFOLD is the introduction of a control variate for all clients, *and* for the server. This control variate consists of a set of values with the same structure as the set of parameters (i.e. it has one value per parameter). Intuitively, it denotes the direction (and magnitude) of the local update of said parameter. If this direction is different from many other clients, the control variate is used to compensate for that difference, which decreases the aforementioned drift.

The general setup of SCAFFOLD is similar to that of federated averaging. The algorithm consists of rounds, which consists of a global part at the server, and a local part which happens at all clients simultaneously. at the beginning of each round, the server sends the global model to all clients, as well as its own control variate c. Now, each client makes a pass over its local data to calculate the gradient over its loss function. This also happens in federated averaging, as part of the stochastic gradient descent step. After the gradient has been determined, the parameters get updated as follows:

$$W^i_{r+1} = W^g_r - \eta \nabla L(X^i_{train}, y^i_{train}) + c^g_r - c^i_r \tag{25}$$

$c^g_0$ and $c^i_r$ have been initialized as all zero, as described in 1 Note that if both $c^g$ and $c^i$ are all zero, equation 25 becomes standard SGD, and SCAFFOLD becomes federated averaging for the local step. After the model has been updated, each client also needs to update its control variate. This is done according to the following equation:

$$c^i_{r+1} = c^i_r - c^g_r + \frac{1}{\eta_l} W^i_r - W^i_{r+1} \tag{26}$$

Finally, $W^i_{r+1}$ is sent back to the server. Once the server has received $W^i_{r+1}$ for all $i \in N$, it aggregates the local updates into a new iteration of the global model. This update is executed as follows:

$$W^g_{r+1} = W^g_r + \frac{\eta_g}{N} * \sum_{i=1}^{N} W^i_{r+1} - W^i_r \tag{27}$$

Which is equivalent to the global update of federated averaging, if $\eta_g$ is set to 1 (which is the case during all experiments described in this report). Different from federated averaging is the need to update $c^g$ as well, which is done according to the following equation:

$$c_{r+1}^g = c_r^g + \frac{1}{N} * \sum_{i=1}^{N}(c_{r+1}^i - c_r^i) \tag{28}$$

Once this is done, the next round starts by the server sending out the updated model parameters and $c$. A big difference wich federated averaging is that SCAFFOLD is a *stateful* algorithm, with the control variates functioning as some sort of state for clients and server.

### 3 Federated PCA

After some initial testing, it turned out that the high dimensionality of the AML datasets was problematic for model convergence. To reduce the dimensionality, a principal component analysis (PCA) ([8]) was applied, and the first 100 principal components were chosen to represent the samples in a federated way, described in supplementary Algorithm 1. Our implementation is taken from[34] (see pseudocode 1), with the simplification of doing all singular value decompositions at the same time centrally instead of incrementally, as was originally proposed; The incremental approach is more scalable, but was deemed unnecessary for our purpose. Two versions of normalizing were explored, using a global or a local mean and variance, see pseudocode 2. The local normalization was deemed to perform a lot better, as it also serves as a z1-normalization.

**Algorithm 1** Federated PCA

```
input: glob (bool), n_pca (int)
On all clients i: X^i_norm = normalize(X^i, glob) ▷ normalize either using
  global or local mean, see supplementary algorithm 2
On all clients i: U^i, ∑^i, V^i = SVD(X^i_norm)
On all clients i: send U^i, ∑^i to server
On server: US^g = concat(U^1 * ∑^1, ..., U^N * ∑^N)
On server: U^g, ∑^g, V^g = SVD(US^g)
On server: send U^g_{0:n_pca} to all clients
On all clients i: X^i_pca = X^i * U^g_{0:n_pca}
```

**Algorithm 2** Normalization options for fedPCA

**input:** **glob**(bool)
**if** glob == True **then**
  **On Server:** request metadata from all clients $i$
  **On all Clients:** $mean^i = mean(X^i)$, $s^i = length(X^i)$ ▷ *Send back local mean and dataset size*
  **On server:** Collect $mean^i$, $std^i$ and $s^i$ for all i
  **On server:** $S = \sum_{i=1}^{n} s^i$ ▷ *Calculate total sample size*
  **On server:** $mean^g = \frac{1}{S} \sum_{i=1}^{n} s^i * mean^i$ ▷ *Calculate global mean*
  **On server:** send $mean^g$ to all clients,
  **On all Clients:** $\sigma^i = \frac{1}{s^i} \sum_{j=0}^{s^i} (X_j^i - mean^g)^2$ ▷ *calculate partial variance on each client*
  **On all Clients:** send $\sigma^i$ to server
  **On server:** $\sigma^g = \frac{1}{S} \sum_{i=0} N s^i * \sigma^i$ ▷ *calculate global variance*
  **On server:** send $\sigma^g$ to all clients
  **On all Clients** $i$**:** $X_{norm}^i = \frac{X^i - mean^g}{\sqrt{\sigma^g}}$
**else**
  **On all clients** $i$ **do:** $mean^i = mean(X^i)$, $\sigma^i = var(X^i)$ ▷ *calculate local mean and variance*
  **On all clients** $i$ **do:** $X_{norm}^i = \frac{X^i - mean^i}{\sqrt{\sigma^i}}$
**end if**

### 4 Supplementary Tables

**Table 1** Results on the two kinase datasets

| | | KDR | | ABL1 | |
|---|---|---|---|---|---|
| | | Final accuracy (std) | Mean AUC (std) | Final accuracy (std) | Mean AUC (std) |
| LR | Central | 0.75 (0.007) | 0.69 (0.006) | 0.78 (0.01) | 0.75 (0.012) |
| | Federated | 0.74 (0.004) | 0.68 (0.005) | 0.78 (0.01) | 0.74 (0.018) |
| SVM | Central | 0.76 (0.007) | 0.7 (0.004) | 0.78 (0.015) | 0.75 (0.017) |
| | Federated | 0.74 (0.005) | 0.69 (0.007) | 0.77 (0.01) | 0.74 (0.017) |
| FNN | Central | 0.75 (0.002) | 0.68 (0.002) | 0.73 (0.004) | 0.71 (0.0) |
| | Federated | 0.75 (0.005) | 0.68 (0.004) | 0.72 (0.003) | 0.71 (0.0) |
| CNN | Central | 0.72 (0.061) | 0.65 (0.053 | 0.76 (0.014) | 0.74 (0.015) |
| | Federated | 0.78 (0.009) | 0.73 (0.012) | 0.77 (0.005) | 0.76 (0.001) |
| GBDT | Central | 0.82 (0.0) | 0.79 (0.0) | 0.81 (0.001) | 0.79 (0.001) |
| | Federated | 0.73 (0.009) | 0.69 (0.003) | 0.78 (0.011) | 0.75 (0.005) |

**Table 2** the architectures of the FNN

| | Layer 1 | Layer 2 | Layer 3 |
|---|---|---|---|
| MNIST2 | Fully Connected: 784 x 100 | ReLU | Fully Connected: 100 x 2 |
| MNIST4 | Fully Connected: 784 x 100 | ReLU | Fully Connected: 100 x 4 |
| fashion MNIST | Fully Connected: 784 x 100 | ReLU | Fully Connected: 100x10 |
| AML | Fully Connected: 100 x 100 | ReLU | Fully Connected: 100x2 |

**Table 3** the CNN architectures

| | Layer 1 | Layer 2 | Layer 3 |
|---|---|---|---|
| MNIST2 | Convolutional: kernel size:3 stride: 1 padding: 1 | Max pool: kernel = 2 stride = 2 | Fully Connected: 196 x 2 |
| MNIST4 | Convolutional: kernel size:3 stride: 1 padding: 1 | Max pool: kernel = 2 stride = 2 | Fully Connected: 196 x 4 |
| fashion MNIST | Convolutional: kernel size:3 stride: 1 padding: 1 | Max pool: kernel = 2 stride = 2 | Fully Connected: 196 x 10 |
| AML | Convolutional: kernel size:3 stride: 1 padding: 1 | Max pool: kernel = 2 stride = 2 | Fully Connected: 25 x 2 |

**Table 4** performances on the MNIST datasets, fashion MNIST and A2 of the AML dataset

| | | | MNIST2 | | MNIST4 | | fashion MNIST | | A2 | |
|---|---|---|---|---|---|---|---|---|---|---|
| | | | Final accuracy (std) | Mean AUC (std) | Final accuracy (std) | Mean AUC (std) | Final accuracy (std) | Mean AUC (std) | Final accuracy (std) | Mean AUC (std) |
| LR | Central | | 0.97 (0.001) | 0.97 (0.001) | 0.93 (0.001) | 0.91 (0.001) | 0.74 (0.003) | 0.66 (0.005) | 0.98 (0.002) | 0.98 (0.002) |
| | Federated | IID | 0.97 (0.002) | 0.97 (0.002) | 0.93 (0.001) | 0.91 (0.001) | 0.78 (0.003) | 0.66 (0.005) | 0.98 (0.003) | 0.98 (0.001) |
| | | CI | 0.97 (0.001) | 0.96 (0.002) | 0.93 (0.001) | 0.91 (0.001) | 0.74 (0.005) | 0.65 (0.005) | 0.98 (0.002) | 0.98 (0.001) |
| | | SI | 0.97 (0.002) | 0.97 (0.002) | 0.93 (0.002) | 0.91 (0.002) | - | - | 0.98 (0.003) | 0.98 (0.001) |
| SVM | Central | | 0.96 (0.003) | 0.92 (0.005) | 0.89 (0.006) | 0.78 (0.008) | 0.75 (0.003) | 0.67 (0.005) | 0.96 (0.006) | 0.91 (0.001) |
| | Federated | IID | 0.96 (0.003) | 0.92 (0.005) | 0.89 (0.006) | 0.78 (0.008) | 0.75 (0.003) | 0.67 (0.005) | 0.95 (0.003) | 0.91 (0.001) |
| | | CI | 0.96 (0.003) | 0.91 (0.005) | 0.89 (0.005) | 0.78 (0.008) | 0.75 (0.004) | 0.66 (0.005) | 0.95 (0.005) | 0.91 (0.003) |
| | | SI | 0.96 (0.003) | 0.92 (0.005) | 0.89 (0.005) | 0.78 (0.008) | - | - | 0.95 (0.004) | 0.92 (0.002) |
| FNN | Central | | 0.95 (0.000) | 0.90 (0.004) | 0.90 (0.013) | 0.82 (0.006) | 0.67 (0.005) | 0.6 (0.008) | 0.98 (0.000) | 0.97 (0.002) |
| | Federated | IID | 0.95 (0.005) | 0.90 (0.005) | 0.91 (0.008) | 0.84 (0.009) | 0.68 (0.006) | 0.61 (0.004) | 0.98 (0.002) | 0.97 (0.001 |
| | | CI | 0.95 (0.001) | 0.90 (0.005) | 0.91 (0.008) | 0.84 (0.009) | 0.68 (0.006) | 0.61 (0.004) | 0.98 (0.004) | 0.97 (0.003) |
| | | SI | 0.93 (0.010) | 0.90 (0.005) | 0.91 (0.008) | 0.84 (0.009) | - | - | 0.97 (0.002) | 0.98 (0.002) |
| CNN | Central | | 0.94 (0.027) | 0.82 (0.041) | 0.90 (0.013) | 0.75 (0.05) | 0.73 (0.023) | 0.54 (0.075) | 0.87 (0.046) | 0.65 (0.047) |
| | Federated | IID | 0.93 (0.011) | 0.83 (0.030) | 0.92 (0.008) | 0.83 (0.03) | 0.73 (0.021) | 0.56 (0.038) | 0.90 (0.005) | 0.65 (0.031) |
| | | CI | 0.93 (0.013) | 0.83 (0.002) | 0.92 (0.008) | 0.83 (0.03) | 0.73 (0.021) | 0.56 (0.038) | 0.90 (0.005) | 0.65 (0.031) |
| | | SI | 0.93 (0.010) | 0.83 (0.030) | 0.92 (0.008) | 0.83 (0.03) | - | - | 0.90 (0.005) | 0.65 (0.031) |
| GBDT | Central | | 0.98 (0.000) | 0.97 (0.000) | 0.96 (0.0) | 0.94 (0.000) | 0.87 (0.0) | 0.84 (0.0) | 0.98 (0.000) | 0.95 (0.000) |
| | Federated | IID | 0.97 (0.002) | 0.96 (0.001) | 0.95 (0.002) | 0.93 (0.001) | 0.84 (0.001) | 0.83 (0.0) | 0.96 (0.001) | 0.94 (0.001) |
| | | CI | 0.97 (0.002) | 0.94 (0.001) | 0.95 (0.001) | 0.92 (0.001) | 0.23 (0.192) | 0.28 (0.184) | 0.95 (0.002) | 0.92 (0.002) |
| | | SI | 0.97 (0.002) | 0.96 (0.001) | 0.95 (0.001) | 0.93 (0.001) | - | - | 0.97 (0.003) | 0.94 (0.001) |

### 5 Supplementary figures

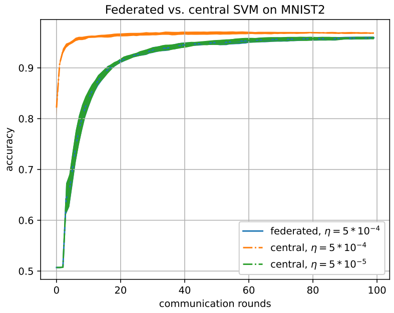


**Fig. 1** the effect of varying learning rates for the comparison between federated and centralized SVM models.

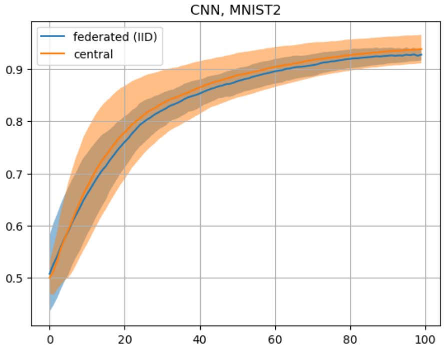


**Fig. 2** Accuracy curves for the centralized and federated CNN on MNIST2 (IID for federated)

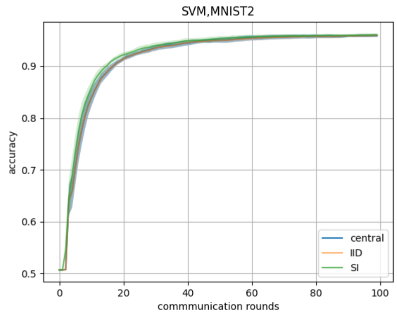


**Fig. 3** Accuracy curves for the federated IID and SI distributions, compared with the central classifier.

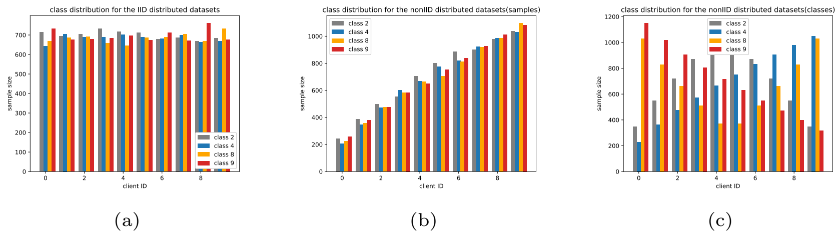


**Fig. 4** Data distributions on MNIST4

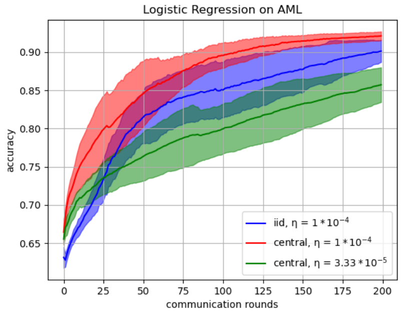


**Fig. 5** LR on the original AML dataset.

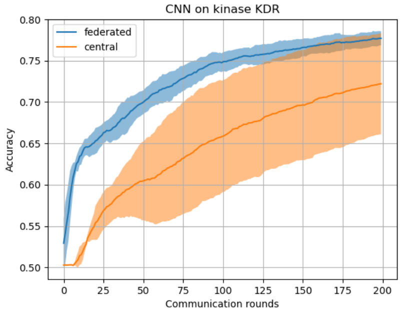


**Fig. 6** CNN on kinase KDR

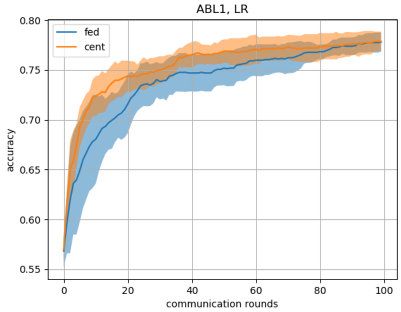


**Fig. 7** LR on kinase ABL1
